## Supplementary figures for "A hierarchy of metabolite exchanges in metabolic models of microbial species and communities"

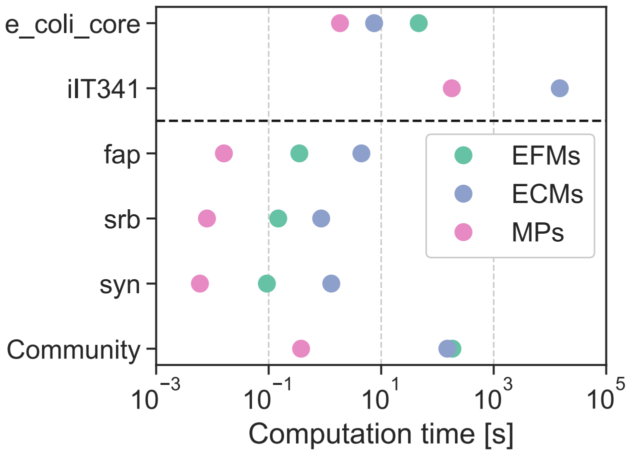


**Fig. S1.** Computation time for enumerating all EFMs, ECMs, or MPs for all models analyzed in this study. Models used to analyze microbial species and models used to analyze a microbial community are separated by a dashed line. EFPs were extracted from EFMs or ECMs and therefore not enumerated separately. All enumerations except ECM enumeration for iIT341 were performed on the same laptop computer (see Methods for details).

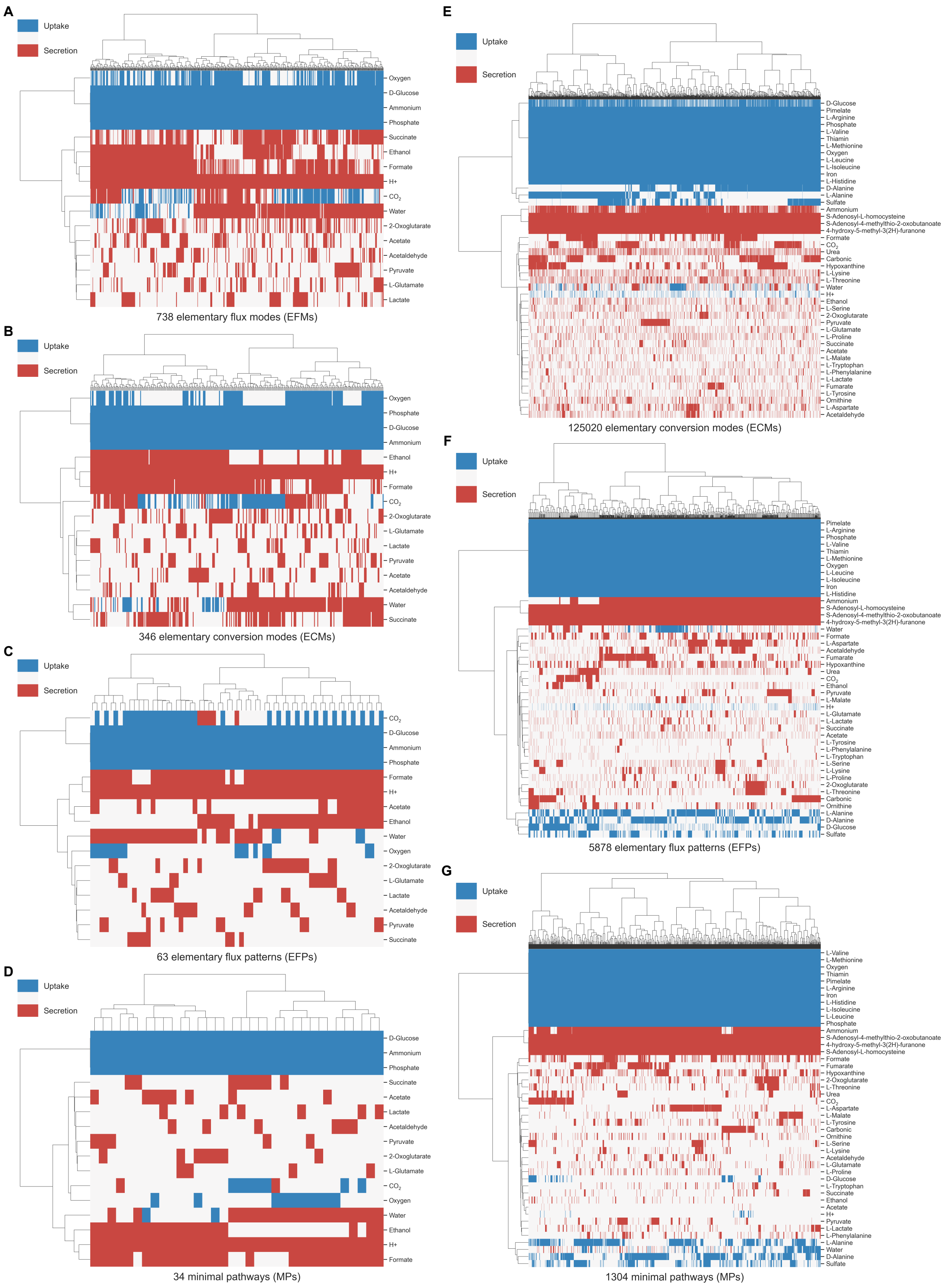


**Fig. S2.** Clustered heatmaps of (**A**) EFMs for e_coli_core, (**B**) ECMs for e_coli_core, (**C**) EFPs for e_coli_core, (**D**) MPs for e_coli_core, (**E**) ECMs for iIT341, (**F**) EFPs for iIT341, and (**G**) MPs for iIT341. Rows are metabolites, columns are pathways, and each cell indicates metabolite uptake (blue) or secretion (red) in a pathway. Rows and columns are clustered by Ward's minimum variance method. Unique growth-supporting flux patterns are shown.

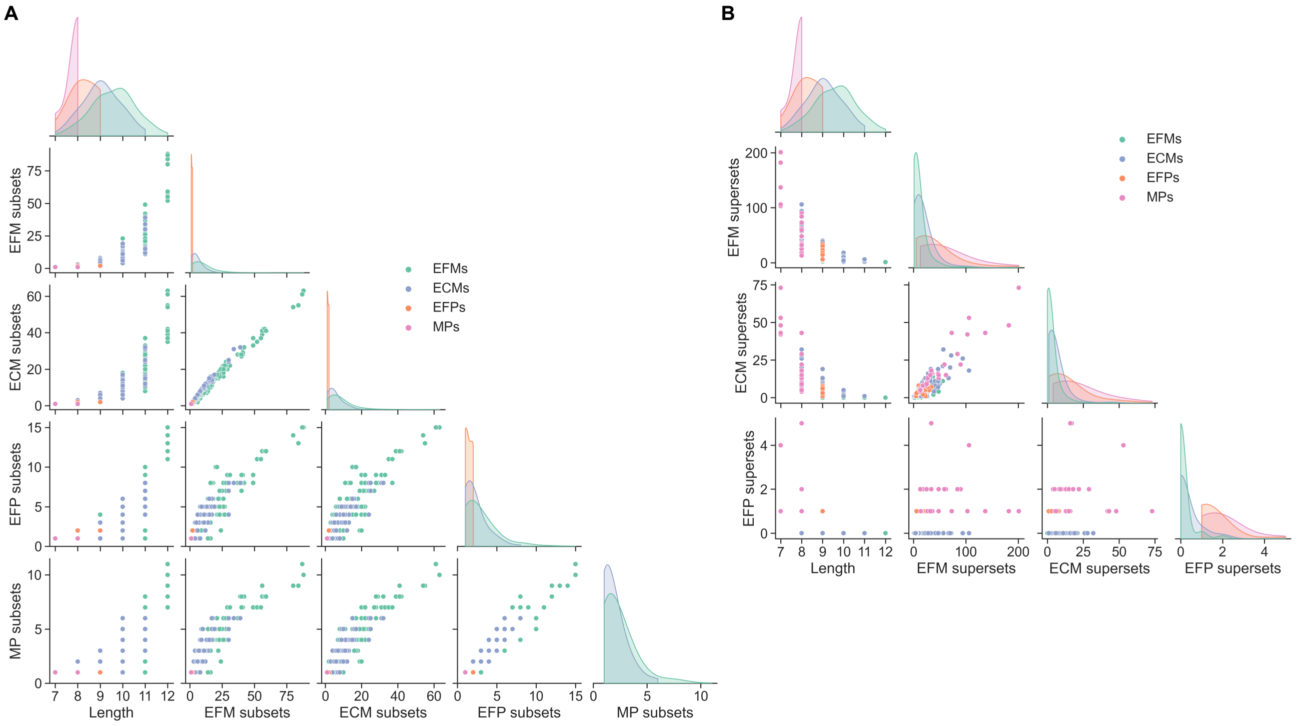


**Fig. S3.** Pairwise relationships between (**A**) pathway length and number of EFM, ECM, EFP, and MP subsets and (**B**) pathway length and number of EFM, ECM, and EFP supersets for e_coli_core.

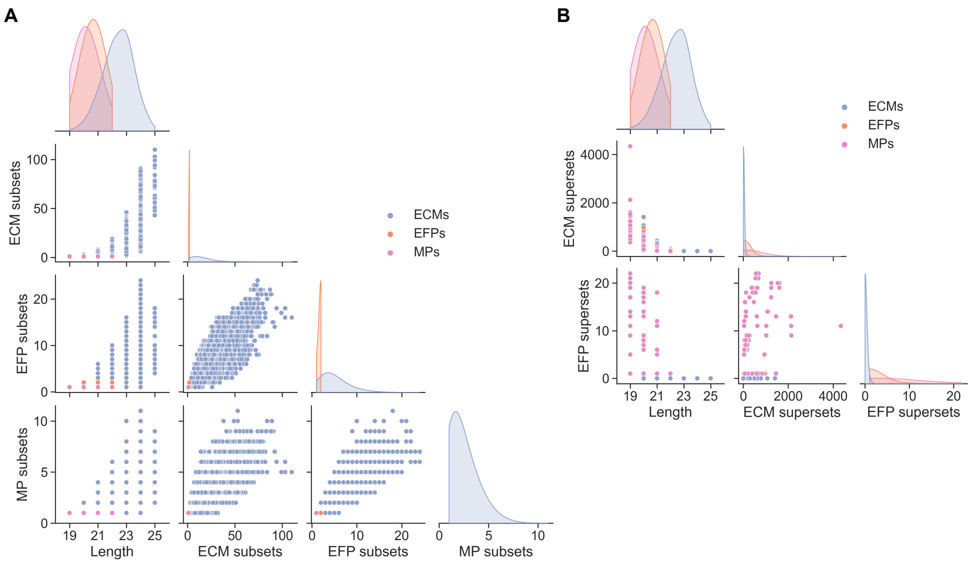


**Fig. S4.** Pairwise relationships between (**A**) pathway length and number of ECM, EFP, and MP subsets and (**B**) pathway length and number of ECM and EFP supersets for iIT341.

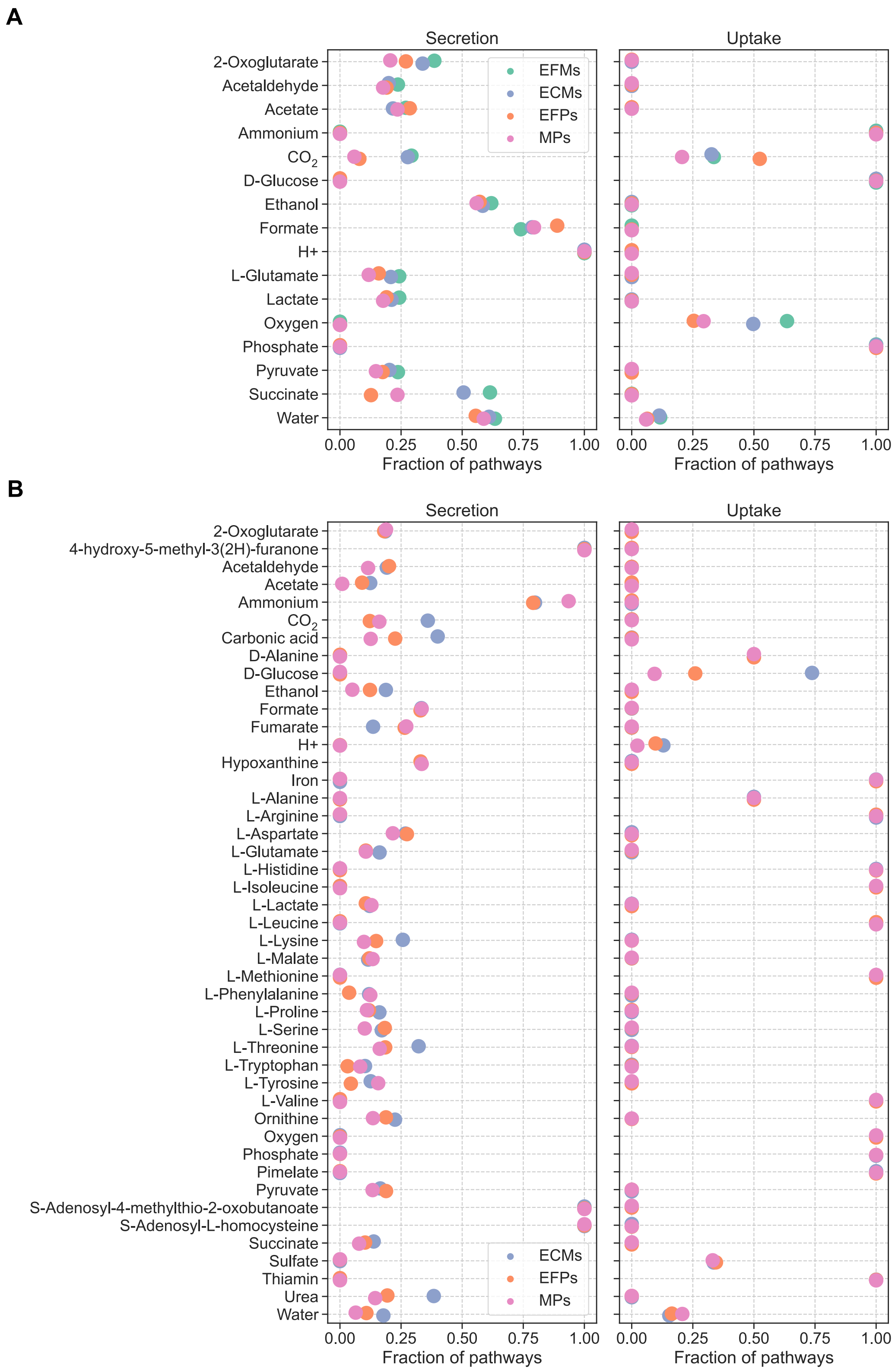


**Fig. S5.** Metabolite exchange frequencies (fraction of pathways including secretion or uptake of each metabolite) for (**A**) e_coli_core and (**B**) iIT341.

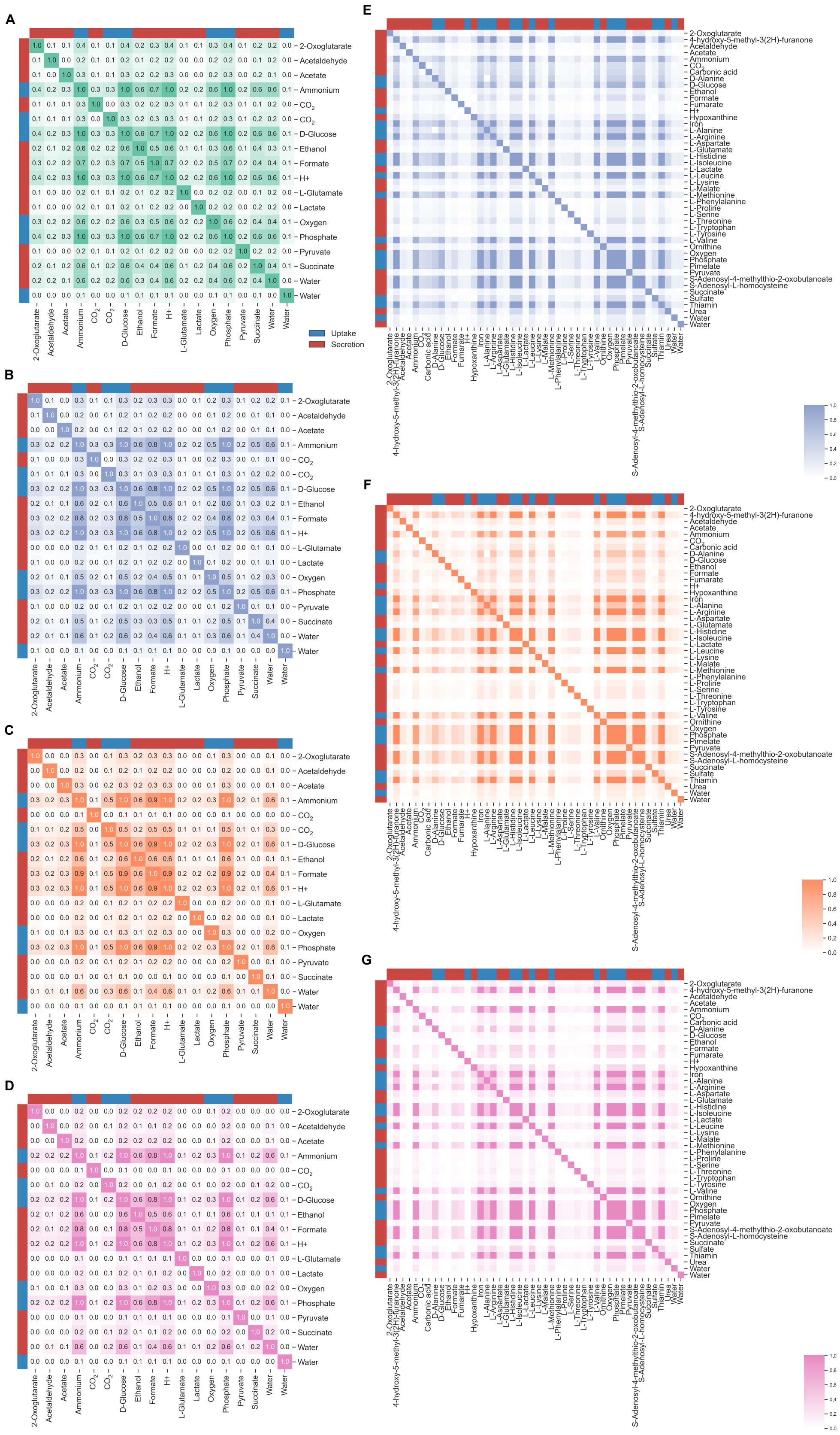


**Fig. S6.** Pairwise metabolite exchange frequencies (fraction of pathways including secretion or uptake of each metabolite pair) for (**A**) EFMs for e_coli_core, (**B**) ECMs for e_coli_core, (**C**) EFPs for e_coli_core, (**D**) MPs for e_coli_core, (**E**) ECMs for iIT341, (**F**) EFPs for iIT341, and (**G**) MPs for iIT341. Rows and columns are metabolites and each cell indicates the pairwise exchange frequency of two metabolites.

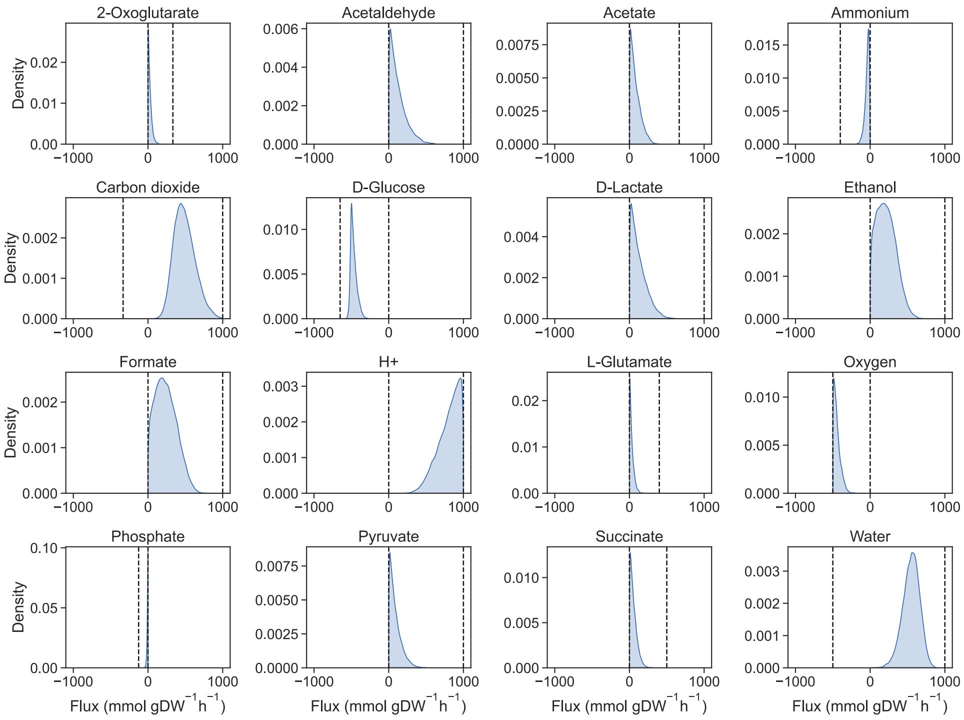


**Fig. S7.** Flux probability distributions for metabolite exchanges in e_coli_core from 100,000 random flux distributions sampled with PTA. Dashed lines indicate feasible flux ranges from FVA.

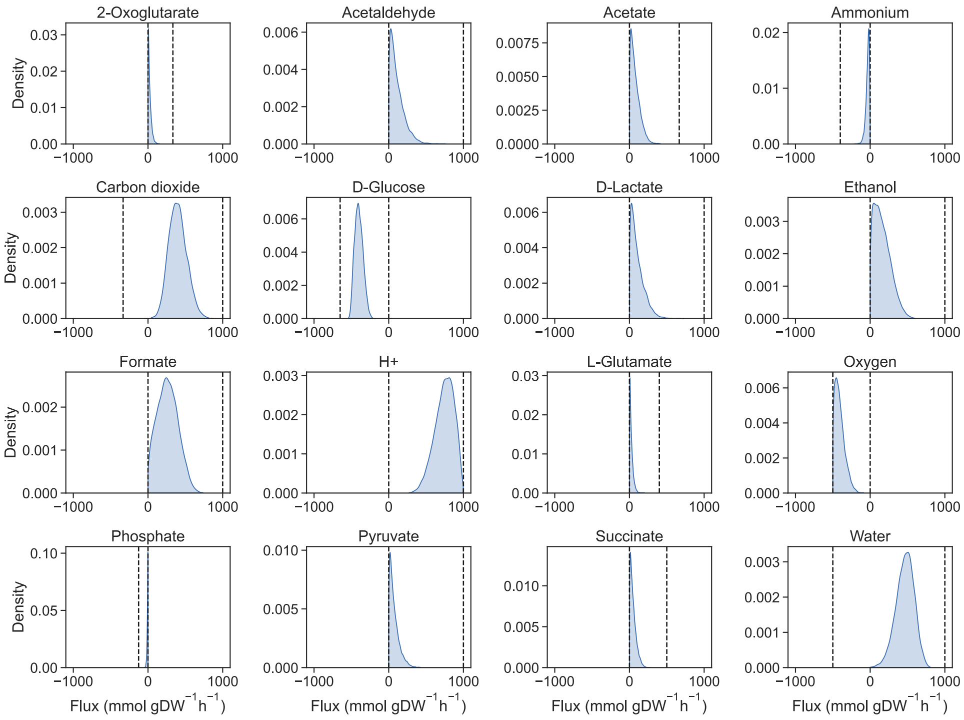


**Fig. S8.** Flux probability distributions for metabolite exchanges in e_coli_core from 100,000 random flux distributions sampled with OptGP. Dashed lines indicate feasible flux ranges from FVA.

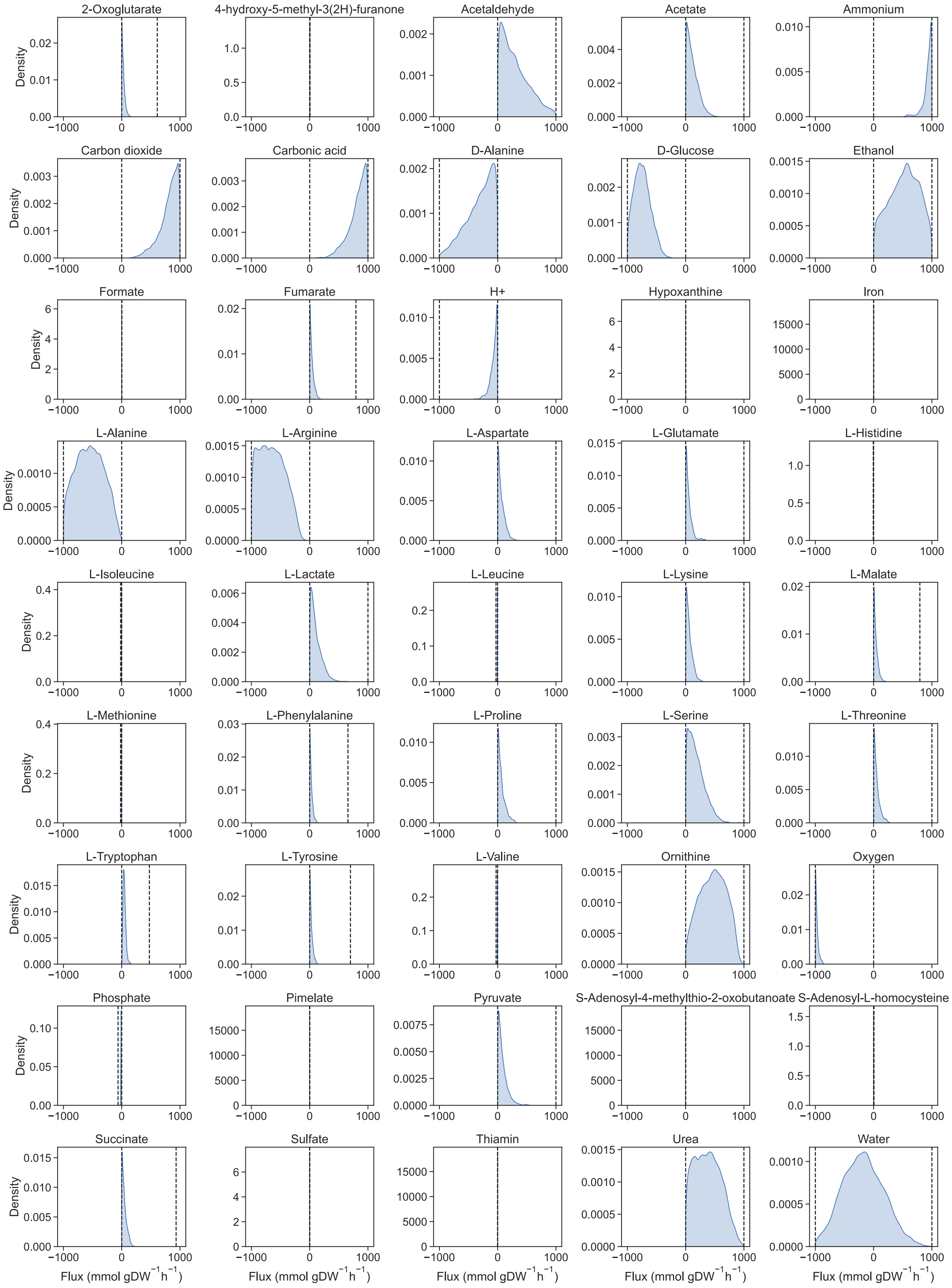


**Fig. S9.** Flux probability distributions for metabolite exchanges in iIT341 from 100,000 random flux distributions sampled with PTA. Dashed lines indicate feasible flux ranges from FVA.

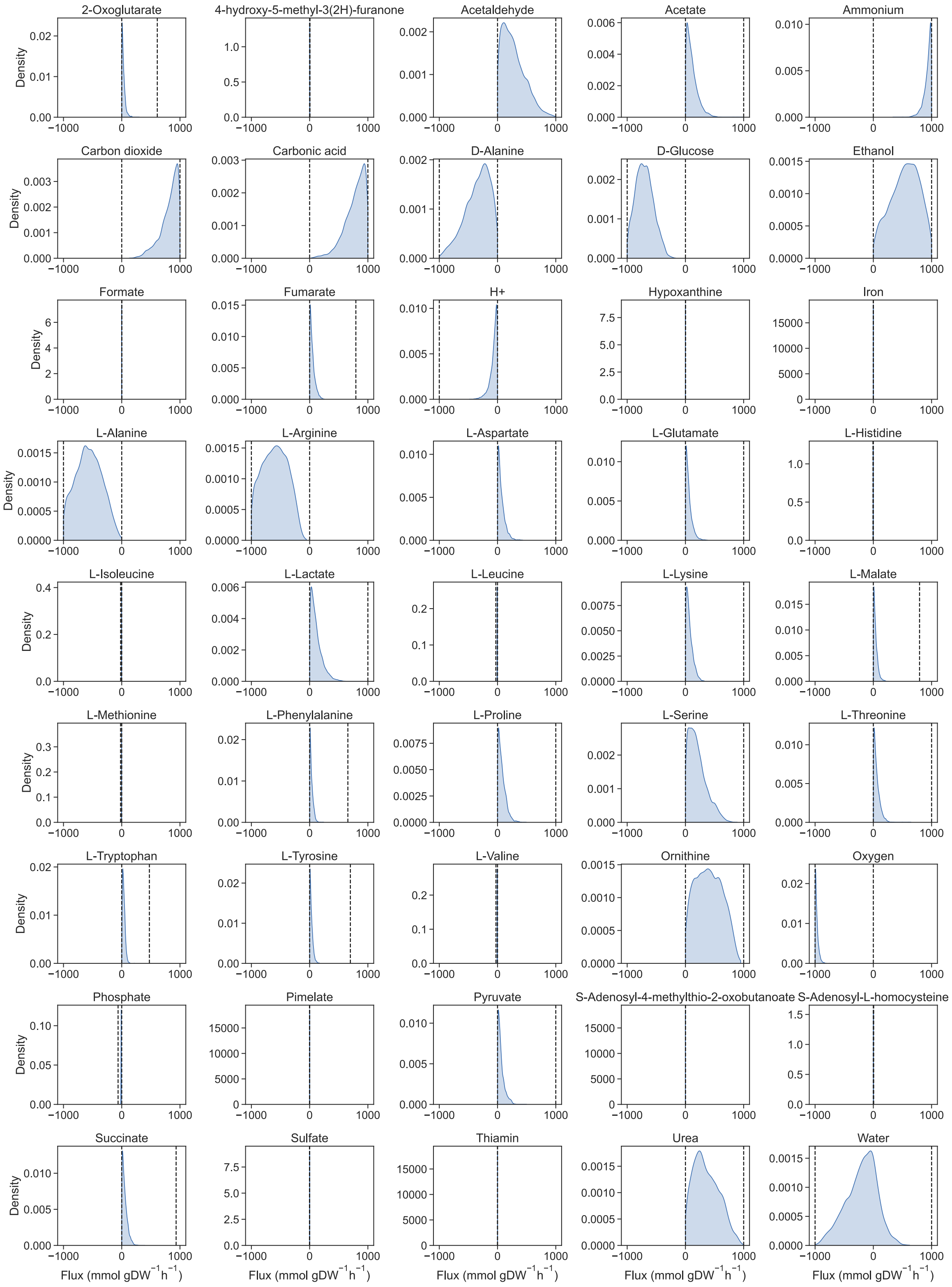


**Fig. S10.** Flux probability distributions for metabolite exchanges in iIT341 from 100,000 random flux distributions sampled with OptGP. Dashed lines indicate feasible flux ranges from FVA.

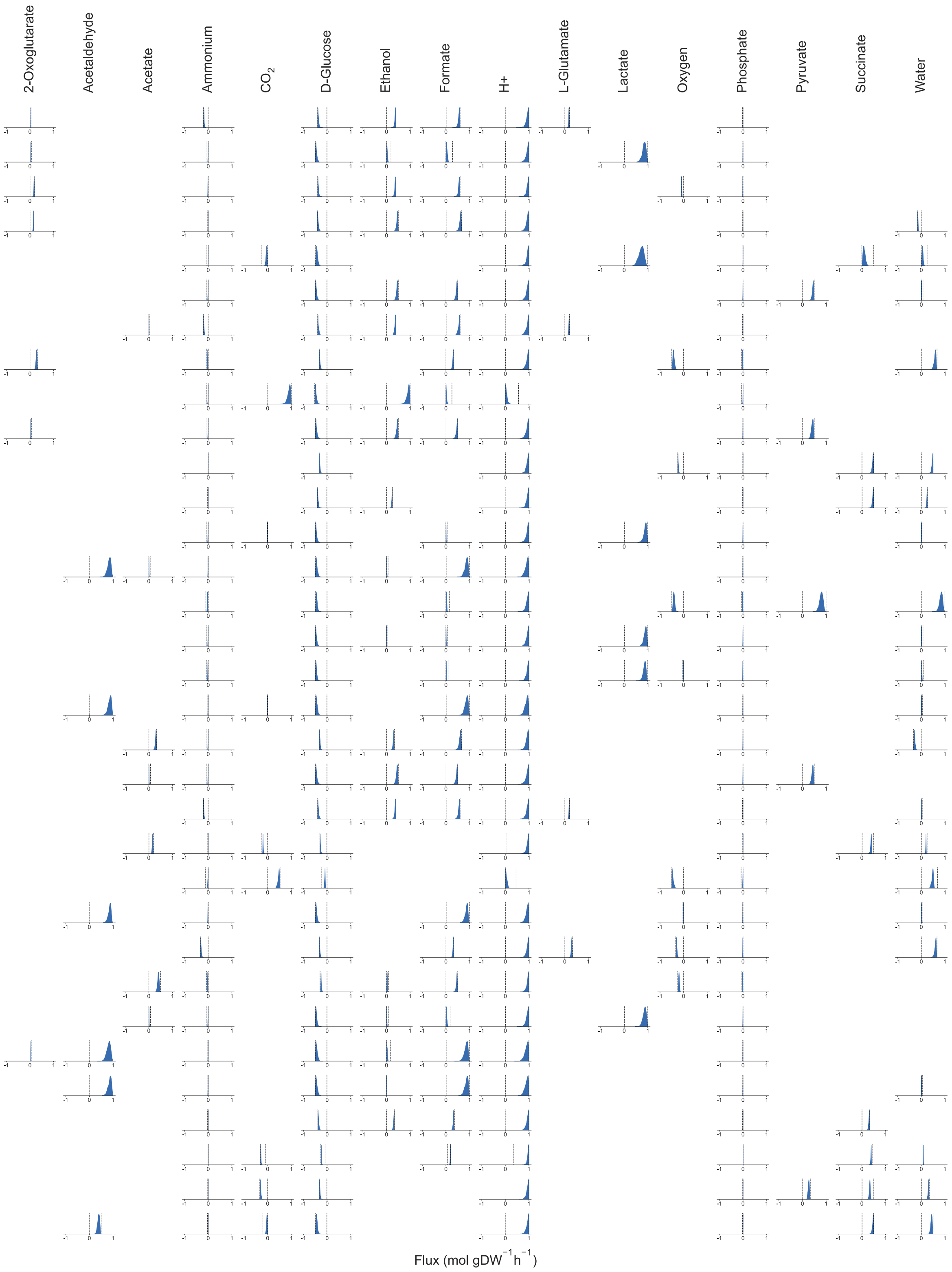


**Fig. S11.** Flux probability distributions for metabolite exchanges in each MP from e_coli_core. OptGP was used to sample 1,000 random flux distributions for each MP. Rows are MPs and columns are metabolites. Dashed lines indicate feasible flux ranges from FVA.

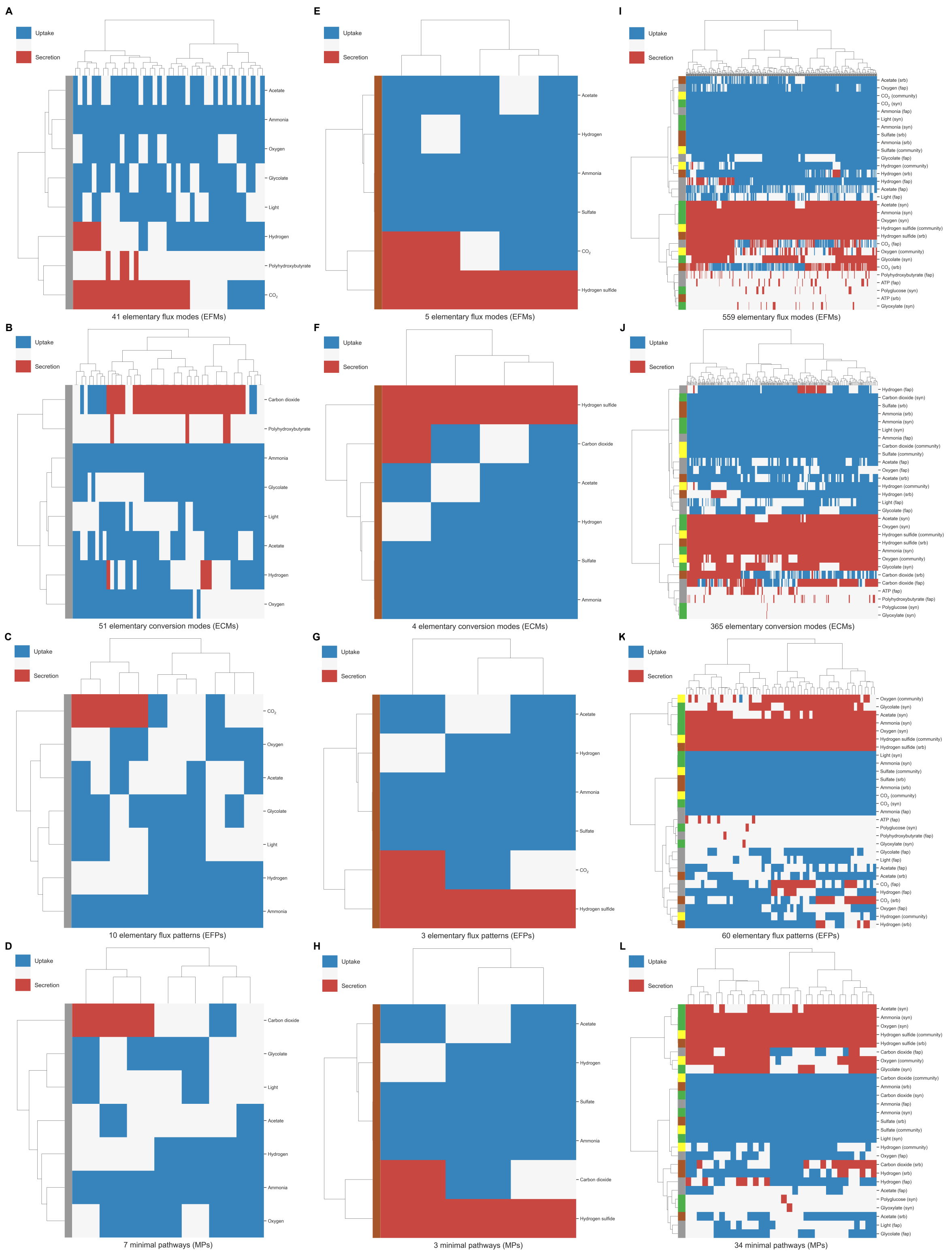


**Fig. S12.** Clustered heatmaps of (**A**) EFMs for fap, (**B**) ECMs for fap, (**C**) EFPs for fap, (**D**) MPs for fap, (**E**) EFMs for srb, (**F**) ECMs for srb, (**G**) EFPs for srb, (**H**) MPs for srb, (**I**) EFMs for syn, (**J**) ECMs for syn, (**K**) EFPs for syn, and (**L**) MPs for syn. Rows are metabolites, columns are pathways, and each cell indicates metabolite uptake (blue) or secretion (red) in a pathway. Rows and columns are clustered by Ward's minimum variance method and rows are colored to indicate individual microbes and the community (yellow). Unique growth-supporting flux patterns are shown.

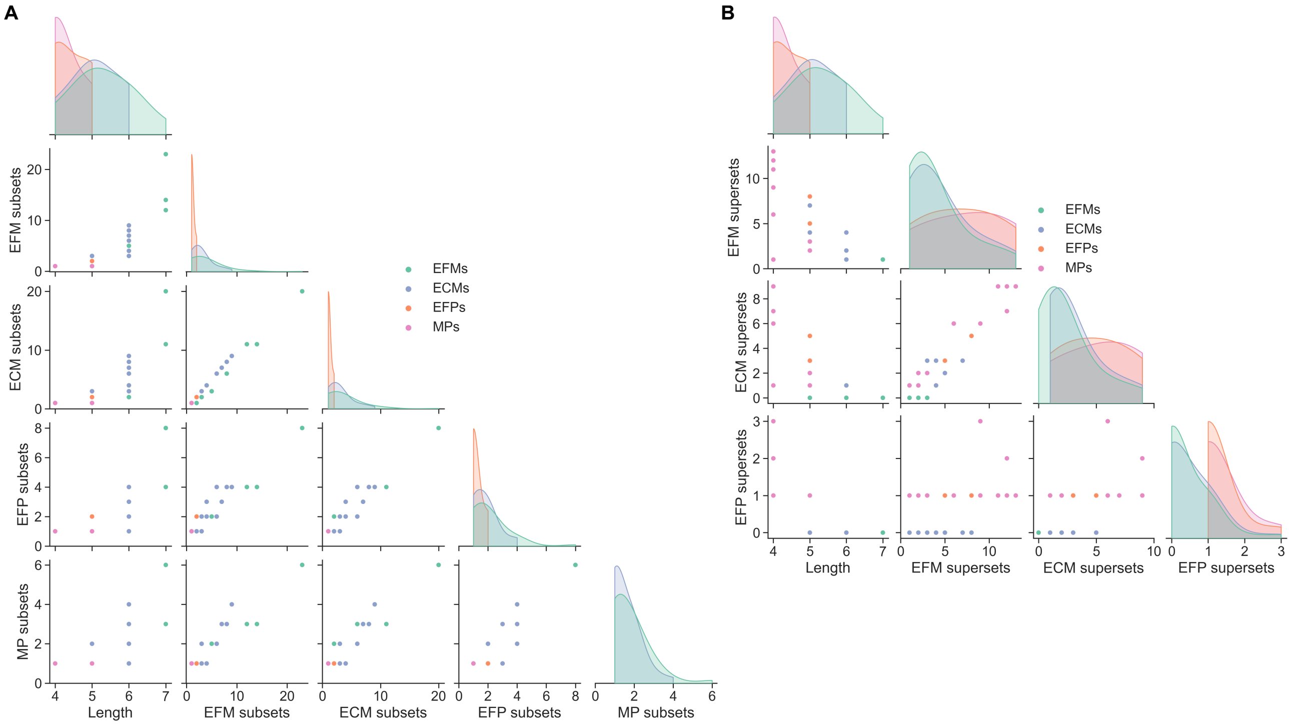


**Fig. S13.** Pairwise relationships between (**A**) pathway length and number of EFM, ECM, EFP, and MP subsets and (**B**) pathway length and number of EFM, ECM, and EFP supersets for fap, srb, and syn individually.

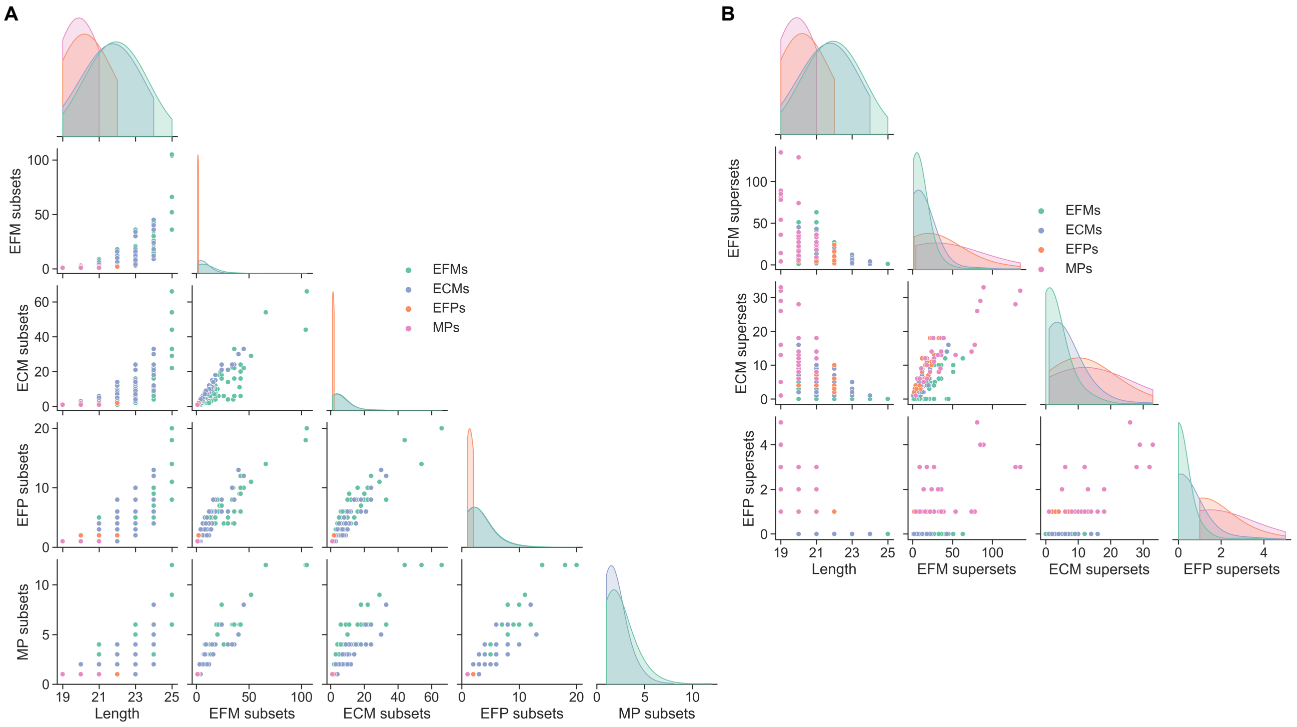


**Fig. S14.** Pairwise relationships between (**A**) pathway length and number of EFM, ECM, EFP, and MP subsets and (**B**) pathway length and number of EFM, ECM, EFP, and MP supersets for the microbial community model.

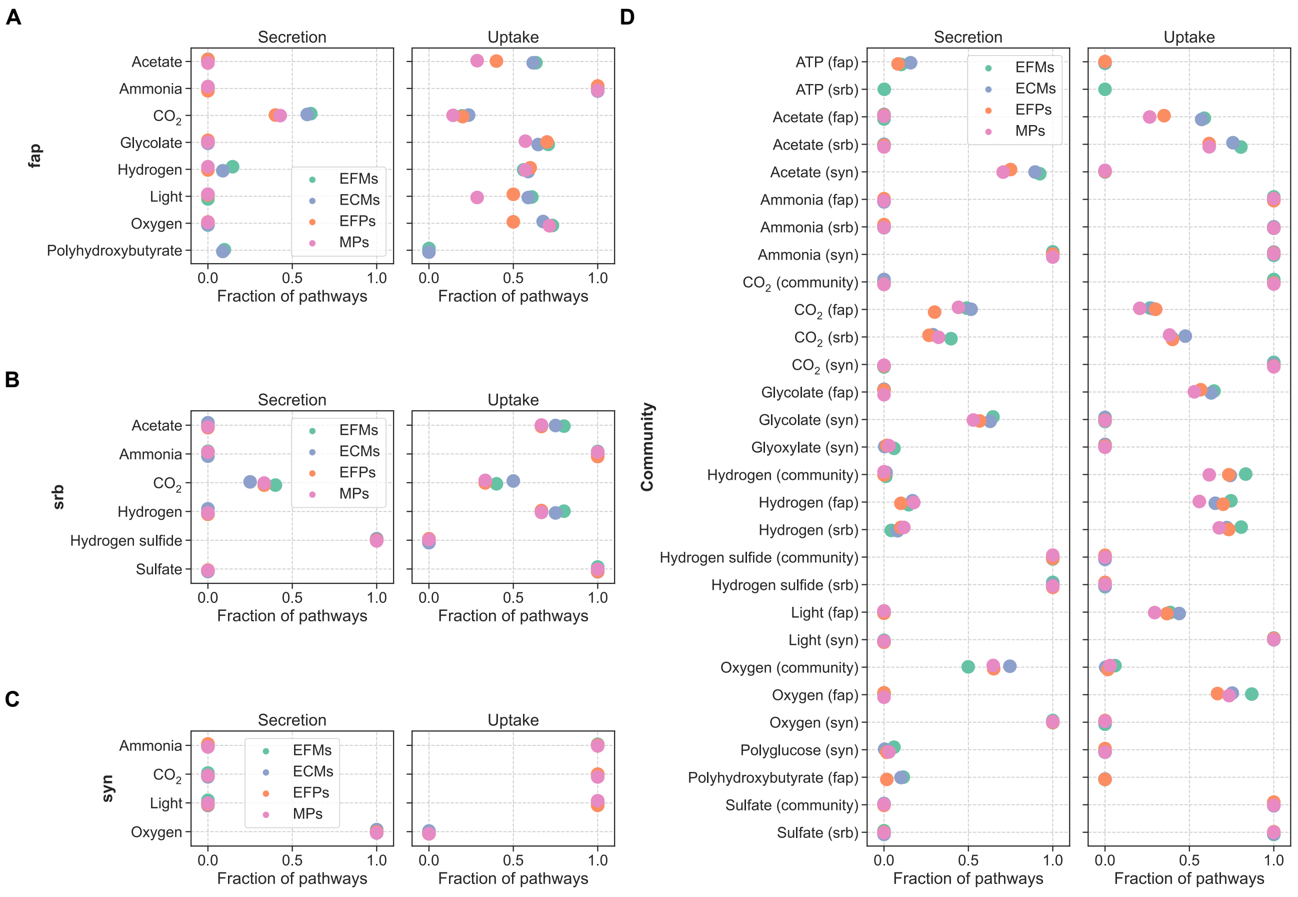


**Fig. S15.** Metabolite exchange frequencies (fraction of pathways including secretion or uptake of each metabolite) for (**A**) fap, (**B**) srb, (**C**) syn, and (**D**) the microbial community model.

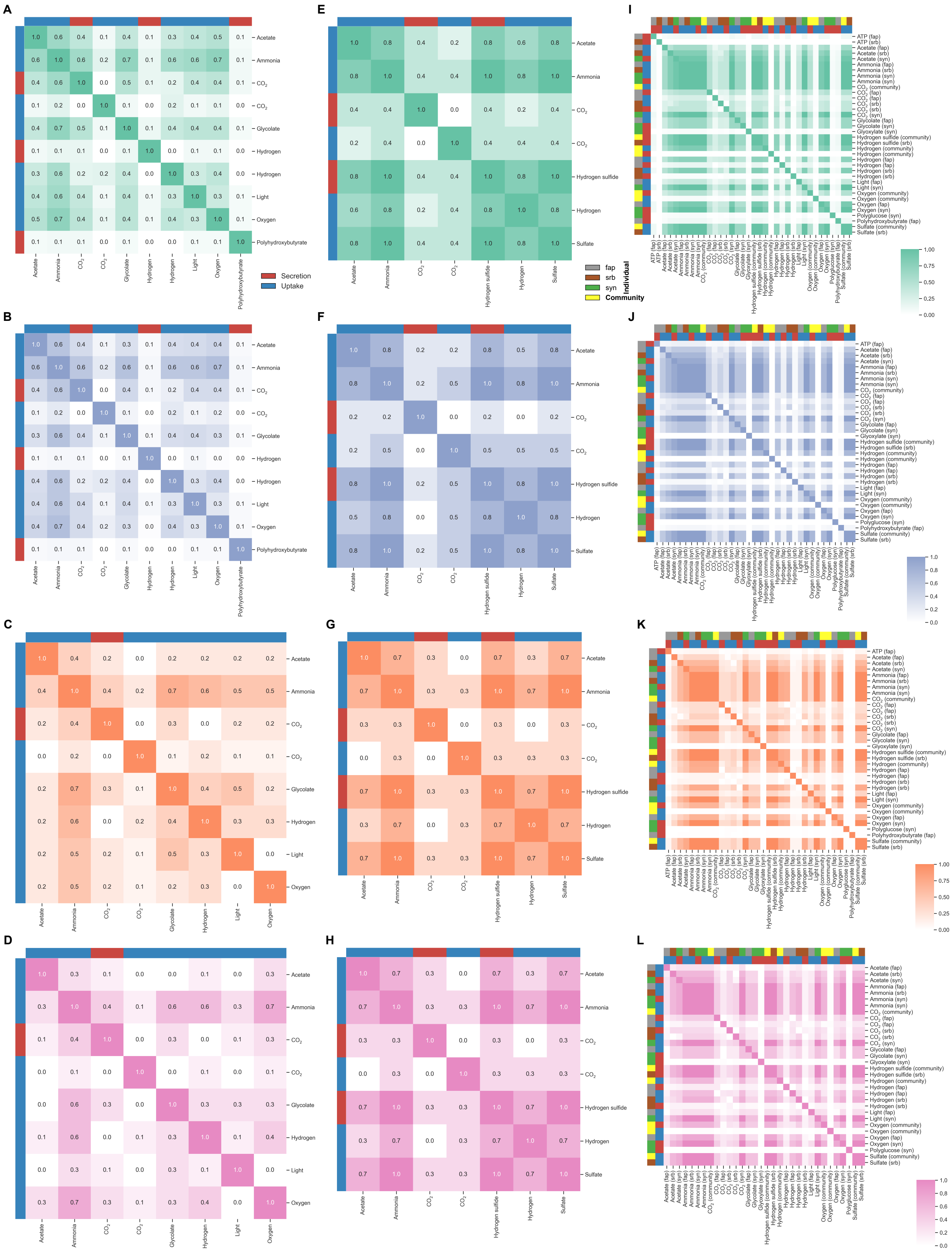


**Fig. S16.** Pairwise metabolite exchange frequencies (fraction of pathways including secretion or uptake of each metabolite pair) for (**A**) EFPs for e_coli_core, (**B**) ECMs for e_coli_core, (**C**) MPs for e_coli_core, (**D**) MPs for iIT341, and (**E**) MPs for iIT341. Rows and columns are metabolites and each cell indicates the pairwise exchange frequency of two metabolites.

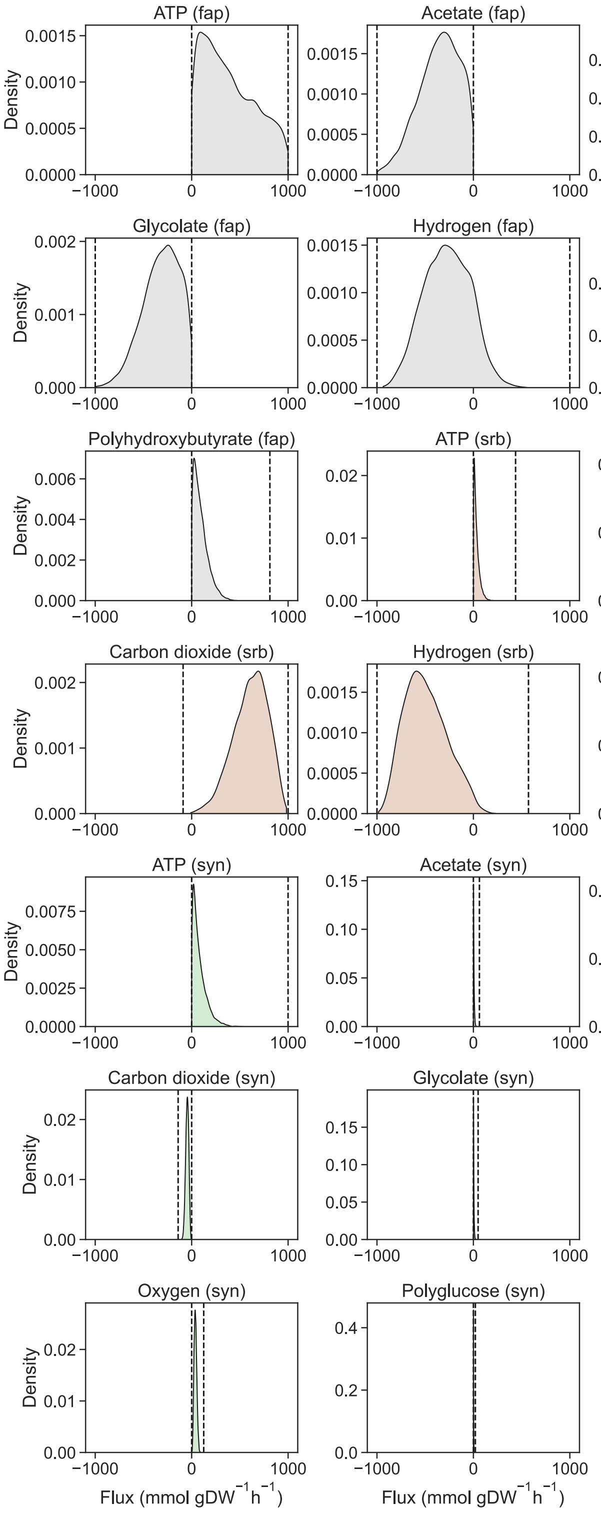


**Fig. S17.** Flux probability distributions for metabolite exchanges in fap, srb, and syn individually from 100,000 random flux distributions sampled with OptGP. Dashed lines indicate feasible flux ranges from FVA.

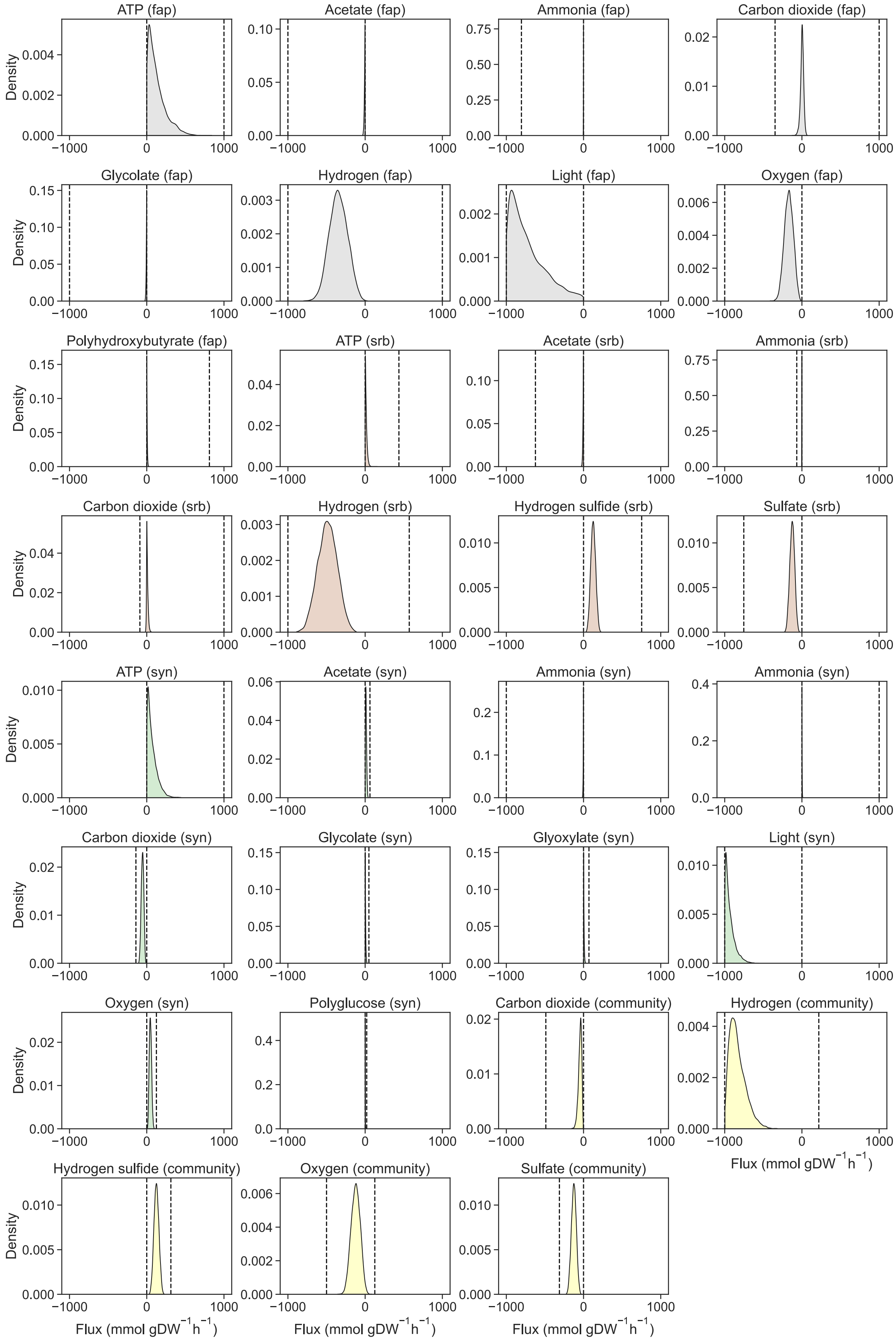


**Fig. S18.** Flux probability distributions for metabolite exchanges in the microbial community model from 100,000 random flux distributions sampled with OptGP. Dashed lines indicate feasible flux ranges from FVA.
